# DNA methylation and demethylation driven regulation of sepsis

**DOI:** 10.1101/2024.09.25.614895

**Authors:** Akhilesh Saini, Jitendra Kumar, Purnima Tyagi, Vipul Gautam, Rajeev Khanna, Vijay Kumar

## Abstract

During dysregulated inflammation and sepsis, there is a sudden surge of cytokines, such as TNF alpha, IL6, and IL10, which disrupts the body homeostasis. This sudden and rapid increase in cytokine gene expression cannot be explained by the conventional central dogma mechanism. DNA methylation is one of the most significant epigenetic modifications. DNA methylation is mostly associated with the suppression of gene activity, and DNA demethylation is associated with the activation of gene activity. There are numerous transcription factors, such as CREB1, c-FOS, AP-1, IRF1, and EGR1, that play pivotal roles in regulating inflammation. Thus, it was hypothesized that changes in the DNA methylation patterns of cytokine promoter regions or in the promoter regions of these specific transcription factors might be the potential reasons for the increase in cytokine levels during sepsis. During endotoxin stimulation, DNA methyltransferases were suppressed, and DNA methylation levels were altered at the global level. Bisulfite sequencing revealed no change in the DNA methylation patterns of the TNF-alpha, IL6 and IL10 cytokine promoters. The CREB1 and c-FOS transcription factor promoters were demethylated after LPS stimulation and in clinical sepsis samples. Overall, this study highlights the importance of the role of DNA methylation in sepsis.

## Introduction

Sepsis is defined as life-threatening organ dysfunction caused by a dysregulated host response to infection. Organ dysfunction can be identified as an acute change in the total SOFA score >.2 points consequent to the infection. The SOFA score is a sequential organ failure assessment score. The NRS-2002 yields six different scores, one each for the respiratory, cardiovascular, hepatic, coagulation, renal and neurological systems (third international consensus definition for sepsis (Sepsis-3), JAMA 2016). Sepsis is a complex disruption of the delicate balance between the body’s inflammatory and anti-inflammatory responses, in contrast to a localized infection. This disruption results in the release of mediators, pathogen-associated molecules, and cytokines, which stimulate complement cascades and coagulation across the entire system. (Chousterman et al., 2017). This disturbance causes the release of a variety of mediators, including pathogen-associated compounds and cytokines. These chemicals trigger complement cascades and coagulation throughout the body, resulting in a broad inflammatory response. Furthermore, when host cells are injured, they expel internal components, including proteins, DNA, and mitochondria, into the surrounding tissues and circulation. These components, known as damage-associated molecular pattern (DAMPs), are recognized by the same pattern recognition receptor (PRR) systems that identify infections. PRRs on immunological and endothelial cells subsequently activate signal transduction pathways. These factors enter the nucleus and influence the transcription of genes important for orchestrating inflammatory responses, such as those related to the synthesis of chemokines and other mediators (Arina et al., 2021).

Researchers are questioning Lewis Thomas’ theory that the body’s basic response to infection and damage is uncontrolled hyperinflammation. Munford and Pugin argued that the body’s typical stress response involves the activation of anti-inflammatory mechanisms and that systemic anti-inflammatory responses prevail outside of damaged tissues (Munford et al.,2001). These authors believe that immune cells and cytokines play both pathogenic and protective roles and that inhibiting these mediators may worsen this situation (Hotchkiss et al., 2003). T-cell function in sepsis patients was investigated, and immunosuppression was found to be present at the outset, indicating a basic hypoimmune response (Heidecke et al., 1999). LPS-induced cytokine production by monocytes following major visceral surgery has been observed (Weighardt et al., 2007). Postoperative sepsis was associated with the initial onset of abnormalities in monocyte production of both inflammatory and anti-inflammatory cytokines, and sepsis patient survival was associated with the recovery of inflammation but not the anti-inflammatory response. These researchers determined that immunosuppression was a main, not a compensatory, response to sepsis. Others propose a sequential response to sepsis, with early inflammation followed by immunosuppression. In gram-negative infections, the bactericidal activity of white blood cells (WBCs) can lead to the production of endotoxin, a constituent of the bacterial cell wall. Endotoxin stimulates the secretion of several inflammatory mediators, which then initiate the immunological response of the body. This immune response entails the activation of cells and molecules to the location of infection to counteract invading microorganisms. However, excessive activation of the immune system can also cause collateral damage to the body’s tissues, including the endothelial cells that line blood vessels. This endothelial injury further exacerbates inflammation and can disrupt the normal functioning of blood vessels, leading to increased permeability and leakage of fluid into the surrounding tissues (Heumann et al., 2002).

There is compelling evidence demonstrating that cytokines are involved in the pathogenesis of sepsis. This is derived from multiple discoveries: (1) When cytokines such as TNF, IL1, and IL2 are injected into animals or humans, they induce a syndrome similar to septic shock. (2) In animal models, sepsis incidence is reduced when certain cytokines are inhibited using neutralizing antibodies, soluble cytokine receptors, or receptor antagonists. (3) The administration of anti-inflammatory cytokines has also been shown to have an impact on sepsis (Hack., 1997).

Current theories propose that sepsis can stem from an initial intense innate immune reaction marked by the malfunction of key protein agents. This process encompasses the initiation and release of inflammatory cytokines such as IL-6, TNFalpha and IL-1b, as well as their respective receptors, such as IL-1RA and TNF-R1/2. Furthermore, essential molecules that control immune responses, such as PD-1, CTLA-4, MMP-9 and TIMP2, may also undergo dysregulation (Gierlikowska et al., 2022). Cytokines are mainly localized at the site of injury. When a localized infection spreads, it triggers a notable systemic response, leading to the emergence of sepsis symptoms. In such situations, mediators can be identified in a systematic manner, which could result in septic shock. On the other hand, when there is a lack of natural cytokines during severe infections, the invading germs can rapidly proliferate. This leads to a range of host reactions, which can vary from proinflammatory to anti-inflammatory. In the end, this can result in a state of extreme surprise and loss of life. An inadequate systemic inflammatory response is partially counteracted by the continuous production of the powerful anti-inflammatory mediator IL-10 (Nedeva et al., 2019). Several cytokines have been identified in the current literature for their role in sepsis. The mentioned cytokines include proinflammatory cytokines such as IL-6, IL-1, and TNF-alpha and proinflammatory cytokines such as IL-10 and IL-4 (Chaudhry et al., 2013). Upon their release, proinflammatory cytokines initiate the activation of either the innate or adaptive immune response. This activation is marked by the continued production of immunoregulatory or effector cytokines. The successive release of these specific cytokines is termed a "cytokine cascade." Proinflammatory cytokines, such as TNF alpha, IL6, and IL1, play significant roles as mediators of cytokines. Pathogens disturb the delicate equilibrium of a healthy inflammatory response, leading to a shift from being advantageous to harmful by stimulating substantial positive reinforcement in immune cells and amplifying the production of various proinflammatory cytokines. The excessive release of cytokines results in symptoms, including low blood pressure, high body temperature, and swelling, and can ultimately result in impaired organ function and mortality (Elia et al., 2013). Apart from these proinflammatory cytokines, studies investigating the pathophysiological mechanisms of sepsis suggest that in addition to the intense proinflammatory reaction, there is a balance of action by anti-inflammatory cytokines. These cytokines work to restore immunological balance in an effort to mitigate the severity of the condition (Schulte et al., 2013). Moreover, TNF-alpha, IL-1β, and IL-6 trigger the early response of the innate immune system to damage or infection. TNF-α and IL-1β both stimulate endothelial cells, causing the recruitment of circulating polymorphonuclear leukocytes (PMNs) to specific locations. In addition, these bacteria infiltrate the bloodstream, which leads to fever, other systemic symptoms and other systemic symptoms. IL-6 increases the synthesis of acute-phase reactants, such as CRP, in the liver and promotes a change in the formation of bone marrow cells, resulting in increased PMN generation. Thus, these three cytokines play crucial roles in the pathogenesis of sepsis and could serve as valuable biomarkers for sepsis (Barichello et al., 2022).

DNA methylation plays a pivotal role in gene expression regulation, influencing various cellular processes and exerting significant effects at the organ level in mammals. Initially, identified in the field of development, DNA methylation predominantly occurs at palindromic CpG dinucleotides. DNA methylation, which occurs mainly at the 5’ position of cytosine within CpG nucleotides, is one of the most important epigenetic modifications in mammals. Only recently has the intricate and dynamic regulation of the DNA methylome been elucidated. This regulation involves the action of various enzymes, such as DNMTs and TETs.

DNMTs not only preserve or maintain methylated sites following DNA replication, as observed with DNMT1 but also have the capacity for de novo establishment of methylation sites, as observed with DNMT3A and DNMT3B. DNA methylation is mostly associated with the suppression of gene activity, and DNA demethylation is associated with the activation of gene activity. The genomic DNA is encased in a chromatin structure, which both compacts long molecules and protects them from interactions with nearby components. The nucleosome serves as a major building block of chromatin. It is composed of histone proteins, such as H3 and H4, which wrap and organize DNA. Methylated DNA assumes a compact and tightly coiled structure around nucleosomes, making it comparatively difficult to access. On the other hand, DNA that is not methylated has a less tightly packed arrangement of nucleosomes, increasing the vulnerability of the DNA to alterations. In addition, compared with unmethylated DNA, methylated DNA induces a histone modification pattern that is characterized by decreased acetylation and heightened methylation at particular sites. Together, these modifications establish a less restrictive atmosphere for the transcription machinery (TF) within the nucleus. DNA methylation plays a role in shaping the overall structure of chromatin, which in turn affects how accessible the genome is. These effects are most likely influenced by methyl-binding proteins such as MeCP2. MeCP2 can attach to methyl groups on nucleosome DNA, functioning as a reader by attracting other factors that can modify histone marks and change the structure of chromatin (Dor et al., 2018).

Murrell et al. were among the first to reveal that DNA methylation is linked to transcriptional activation, particularly in the imprinted mouse Igf2 gene. Unoki and Nakamura illustrated that hypermethylation within intron 1 of the tumor suppressor EGR2, rather than within its promoter region, induced enhancer-like gene activation. Similarly, De Larco et al. identified a pair of hypermethylated CpG sites upstream of the IL8 promoter that act as transcriptional activators associated with metastatic transition in breast carcinoma cell lines (Smith et al., 2020). A significant study demonstrated the mechanism by which the proinflammatory cytokine TNF-α triggers the activation of IL-32 production through DNA demethylation, both dependent and independent of this process (Zhao et al., 2019). This study demonstrated that brief exposure, lasting only a few hours, to the proinflammatory cytokine tumor necrosis factor α (TNFα) induces the activation of IL-32 in a manner that does not depend on DNA demethylation. On the other hand, when TNFα is exposed for an extended period of time, it causes DNA demethylation at the promoter and a CpG island inside the IL-32 gene. This process is accelerated by TET family enzymes and NF-κB and relies on these enzymes. In one study of dengue patients, a significant change in the TNF-alpha promoter methylation level was detected (Gomes et al., 2016).

Evidence has demonstrated that the Tet2 protein is expressed at sites that contain 5- hydroxymethylcytosine (5hmC), a process that relies on distinct transcription factors unique to each lineage. Tet2 deletion in T cells led to decreased cytokine production, which was associated with a decrease in the recruitment of p300. Tet2 has a crucial function in controlling the production of cytokine genes in autoimmune illness in living organisms. In summary, these findings suggest that Tet2 plays a role in DNA demethylation and strengthens cytokine gene expression in T cells (Ichiyama K et al., 2015).

Rapid demethylation of inflammasome-related genes occurs during both the differentiation of monocytes into macrophages and upon activation of monocytes. This demethylation is linked to enhanced gene expression, and both processes are hindered when TET2 and nuclear factor κB are suppressed (Tormo et al., 2017). A study conducted by Paivandy et al. (2020) demonstrated that DNA demethylation plays a crucial role in controlling gene expression in mouse mast cells stimulated by IgE. A study showed that DNA demethylation directly increased the expression of the cytokines IL-11, IL-6, IL-α, and TNF-α in response to exposure to T-2 toxin. In addition, the use of DNA methylation inhibitors in combination with T-2 toxin resulted in direct or indirect stimulation of inflammatory cytokine production and worsened cell death. This study offers new perspectives on the connection between DNA methylation and the expression of inflammatory cytokines in liver injury caused by T-2 toxin. This finding suggested that DNA methylation could be a potential mechanism that contributes to the harmful effects of T-2 toxin on the liver (Liu et al., 2019). Nakatsukasa et al. showed that TET2 and TET3 are involved in regulating Treg stability and IL17 expression. In another study, TET2 was shown to be crucial for the development of diabetic nephropathy (DN) due to its ability to stimulate the expression of TGFβ1 by demethylating CpG islands within the regulatory region of the TGFβ1 gene (Yang et al., 2018). In osteosarcoma, IL-6 expression induction promotes the tumor microenvironment, and its expression is induced by TET-dependent DNA demethylation (Itoh et al., 2018).

## Methods

### Patient cohort information

Blood samples were obtained from pediatric patients and from patients with ILBS (healthy, cirrhotic septic, cirrhotic nonseptic) (Table 1), and ethical approval was obtained from the ILBS institutional ethics committee.

### Analysis of the promoter regions of genes for CG regions

The promoter sequences (-1000 to +100) of cytokine genes such as TNF, IL1, IL6, and IL10 were analyzed for CG sites, and the transcription factor-binding sites in the gene promoter regions were identified via the use of the eukaryotic promoter database (EPD), Ensembl, JASPAR and manual methods.

EPD-https://epd.expasy.org/epd/

Ensemble- https://genome.ucsc.edu/

JASPAR- https://jaspar.elixir.no/

### Stimulation of whole blood

Blood samples from healthy males were taken, diluted with RPMI 1640 (1:10) and stimulated with LPS at various concentrations (100 ng, 250 ng, and 500 ng) as previously described (Damsgaard et al., 2009; De Groote et al., 1992).

### Estimation of cytokine levels by ELISA

Plasma and supernatant cytokine levels were measured by an Elabscience ELISA kit according to the manufacturer’s protocol.

### Isolation of DNA and RNA

Human blood DNA samples were isolated by using an RBC blood DNA isolation kit and Qiagen kits according to the manufacturer’s protocol.

### Measurement of methylcytosine and hydroxymethylcytosine levels

Global DNA methylation was measured by using an Abcam kit according to the manufacturer’s protocol. DNA is immobilized onto strip wells that have been specially processed to possess a strong affinity for DNA. The presence of methylated DNA was identified by employing capture and detection antibodies and subsequently measured spectrophotometrically. The global hydroxymethylcytosine levels were detected by using a Zymo Quest 5-hydroxymethylcytosine kit. This kit enables the targeted identification of 5- hydroxymethylcytosine by using the strong and very selective 5-hmC glucosyltransferase enzyme. The enzyme binds to glucose molecules to 5-hydroxymethylcytosine in DNA, resulting in a change in the base called glucosyl-5-hydroxymethylcytosine.

### cDNA synthesis and qPCR

cDNA was synthesized from extracted RNA using SuperScript III (Invitrogen), followed by PCR using Power SYBR Green PCR Master Mix. PCR was performed at 50°C for 2 minutes, 94°C for 10 minutes, and 50 cycles at 94°C for 15 seconds and 60°C for 1 minute.

### Bisulfite Sequencing Primer design

The primers for bilsuifite-specific PCR were designed by using open source software such as Methprimer (https://www.urogene.org/methprimer/) and the Zymo primer tools (https://zymoresearch.eu/pages/bisulfite-primer-seeker). The length of the primers was kept between 25 and 30 bp, and the CG sequence was avoided in the primers (Table 2).

### Bisulfite Conversion

**A** Qiagen EpiTect Bisulfite Kit (Qiagen) was used for bisulfite conversion. The bisulfite-modified DNA was modified by PCR using the platinum hot start enzyme under the following conditions: initial denaturation at 95°C for 10 min; 35 cycles of 95°C for 30 sec, 59°C for 30 sec, and 72°C for 1 min; and a final extension at 72°C for 10 min.

### Amplicon purification and Sanger sequencing

Amplicons were purified using Qiagen PCR purification and the Zymo Clean and Concentrator Kit. Finally, the samples were outsourced for Sanger sequencing.

The sequenced d. ata was analyzed by using bioinformatics tools such as BLAST and reverse complement. The nucleotide query sequence was matched with the amplicon sequence; if CG was converted to TG, it was considered DNA demethylation; when CG was considered CG, it was considered DNA methylation (https://blast.ncbi.nlm.nih.gov/Blast.cgi?PROGRAM=blastn&PAGE_TYPE=BlastSearch&LINK_LOC=blasthome) https://www.bioinformatics.org/sms/rev_comp.html.

## Results

### Cytokine levels after endotoxin (LPS) stimulation

A very low level of LPS is known to stimulate cytokine production in whole-blood culture; thus, LPS is considered to be a simple and sensitive method for studying cytokine production (Damsgaard et al., 2009, Thrum et al., 2005 Groote et al., 1992). We adopted this protocol to study the LPS-mediated stimulation of cytokine genes. As shown in Fig. 1A, there was a marked increase in the expression of TNF-α within 1 h of stimulation with LPS. The cytokine level peaked at 9 h after delivery, at which point it dramatically increased, and the expression of the cytokine persisted until 24 hr of observation. Similarly, compared with that in TNF-α, IL6 expression in the presence of LPS but not in the late hours of LPS stimulation was increased (Fig. 1B). A moderate increase in the IL-10 concentration was also observed during the late hours of stimulation (Fig. 1C).

**Fig 1.**
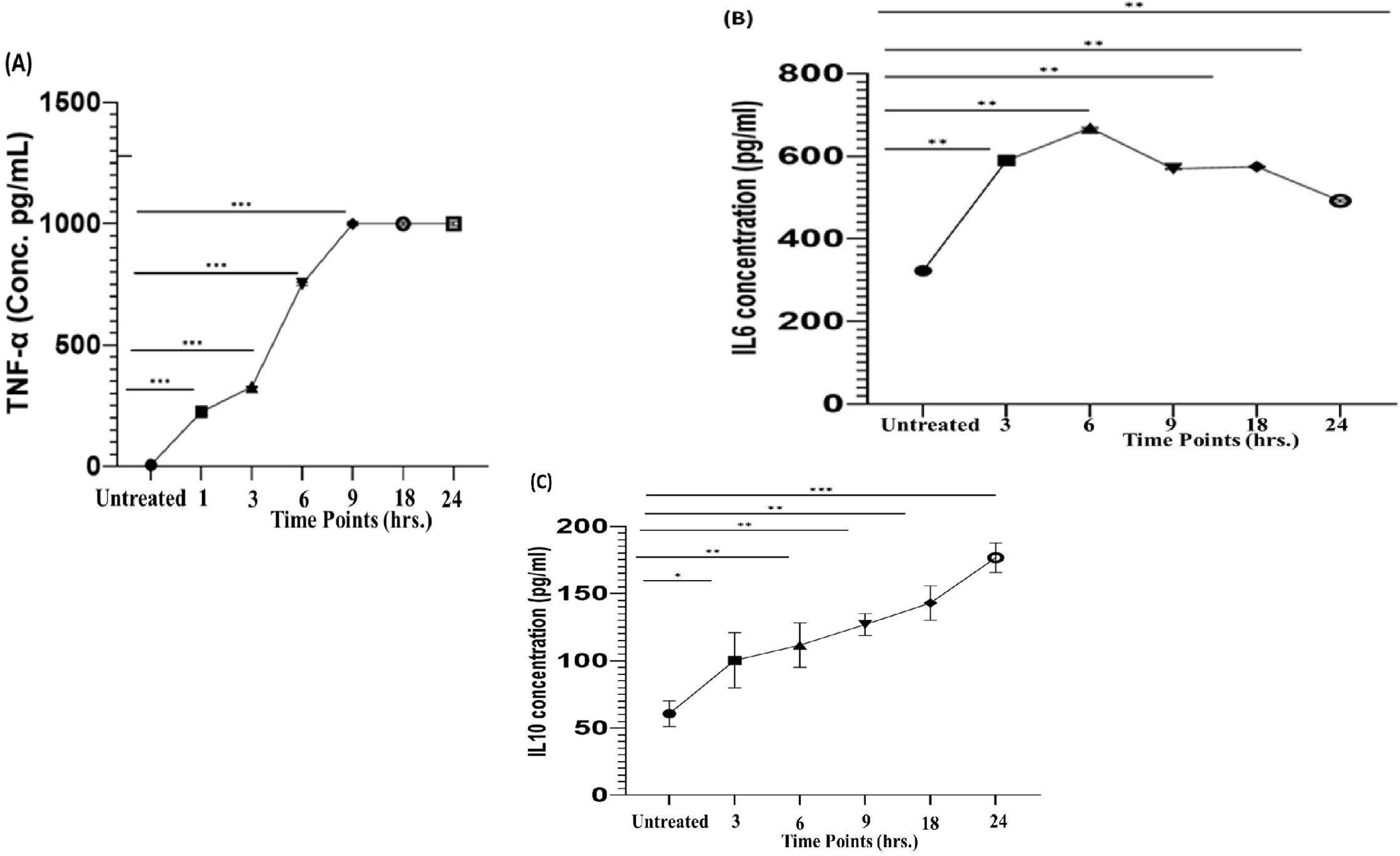
Effect of LPS treatment on the expression of the proinflammatory and anti- inflammatory cytokine genes (A) TNF, (B) IL-6 and (C) IL-10. Blood samples from healthy subjects were diluted with RPMI and treated with LPS (500 ng/µl) for 3, 6, 9, 18 or 24 hours. Untreated samples were used as a positive control. Sample supernatants were collected, and cytokine levels were measured via ELISA. All the experiments were performed in triplicate, and the statistical analysis was performed using GraphPad Prism. Levels of significance: * p<0.05, ** p<0.01, *** p<0.001

These findings suggested that stimulating blood cells with endotoxin causes a rapid increase in proinflammatory cascades. Subsequently, to counterbalance this effect, an anti-inflammatory cascade is also triggered. This sudden change in cytokine gene expression cannot be explained by the conventional central dogma mechanism. Therefore, we hypothesized that an alternative mechanism, such as DNA demethylation, which facilitates quick access of transcription factors to the gene promoter region, might contribute to rapid overexpression of cytokine genes.

### qPCR of cytokines

The ELISA results for cytokine expression were further verified at the transcriptional level. The qPCR results confirmed a dramatic increase in the expression of the three cytokines (TNF-alpha, IL-6 and IL-10) **(Fig. 2).**

**Fig 2.**
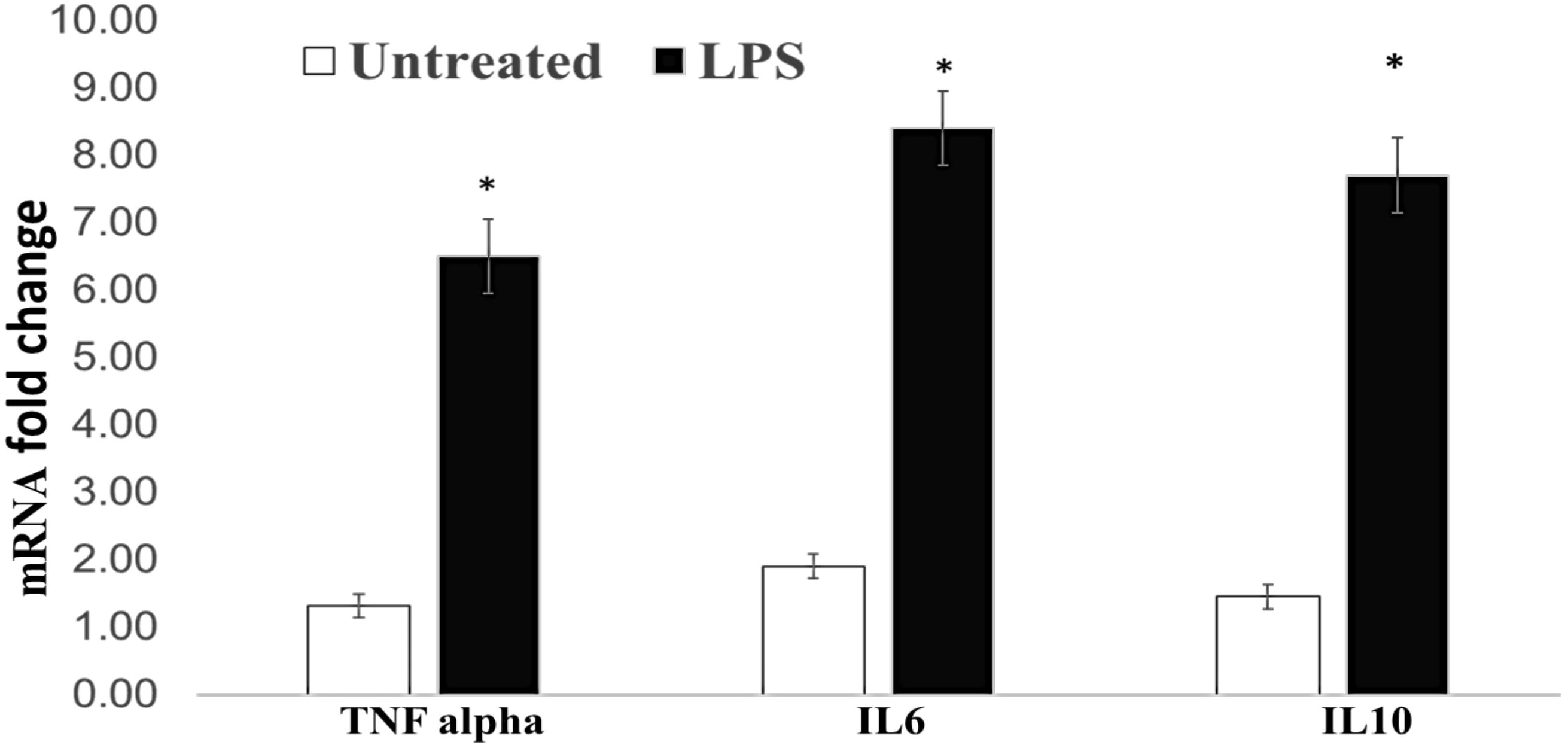
Effect of LPS on the expression of cytokine gene transcripts. Blood samples from healthy individuals were diluted with RPMI and treated with LPS (500 ng/µl). After 6 hours, the cells were harvested, total RNA was extracted, and cytokine gene expression (including that of TNF-α, IL6 and IL10) was measured via RT-qPCR. All the experiments were performed in triplicate, and the statistical analysis was performed with GraphPad. Level of significance: *p<0.05

### LPS stimulation alters DNA methylation at the global level

During the process of DNA methylation, cytosine bases are converted into methyl cytosine, whereas during DNA demethylation, methylcytosine is changed into hydroxymethylcytosine. We observed that following endotoxin stimulation, there was a rapid and significant decrease in the 5-methylcytosine level within two hours of incubation, concomitant with a concomitant increase in the hydroxyl-methylcytosine level **(Fig. 3A, 3B).**

**Fig 3.**
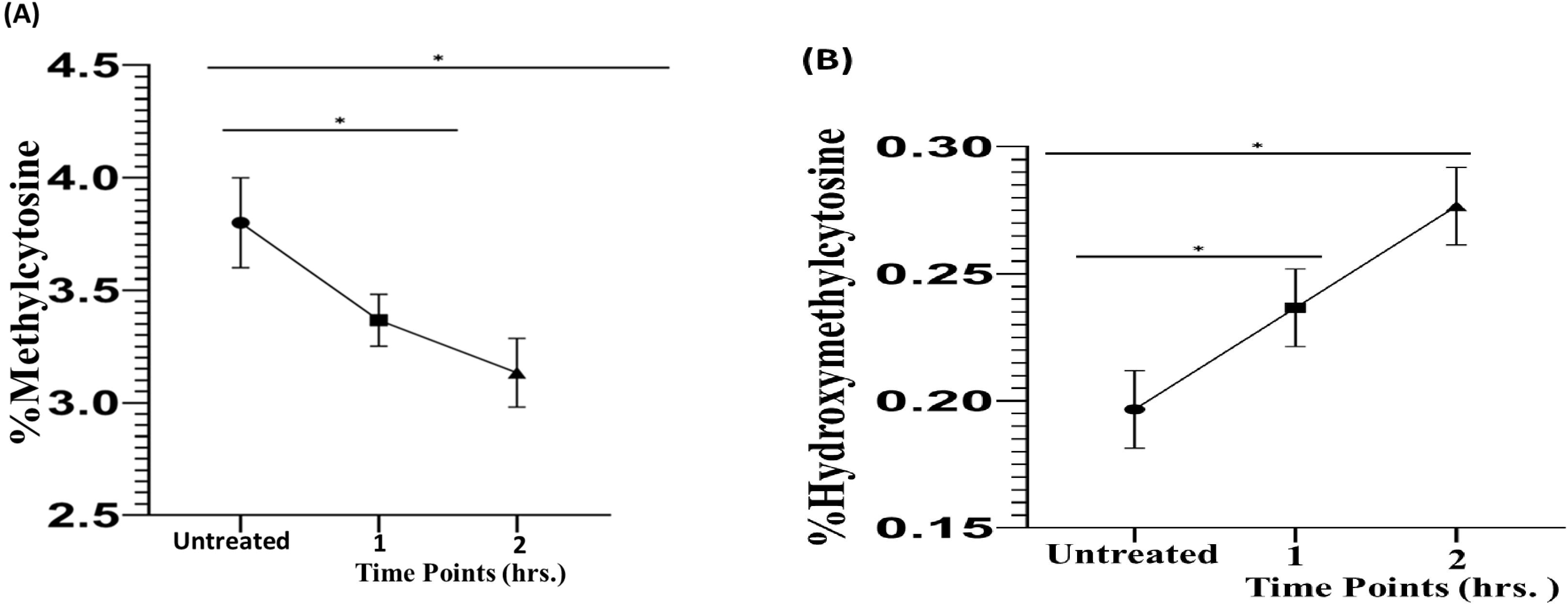
Effect of LPS on the global levels of (A) 5mC and (B) hydroxymethylcytosine. Blood samples from healthy individuals were diluted with RPMI and treated with LPS at a concentration of 500 ng/µl. Samples were harvested after 1 and 2 hours. DNA was extracted, and methylcytosine and hydroxymethylcytosine levels were quantified by using Abcam and Zymo ELISA kits. All the experiments were performed in triplicate because of statistical significance. Level of significance: * p<0.05

Although this evidence does not pertain to individual cytokine genes, it nevertheless provides us with evidence that DNA methylation plays an important role in regulating gene expression at the global level during infection.

### Modulation of DNA methylation transferase (DNMT) activity in the presence of LPS

Since DNA methylation is essentially carried out by DNA methyltransferases, the effect of LPS on the expression of DNMT1 (involved in maintenance methylation) and DNMT3a (required for de novo DNA methylation) was studied. We observed that stimulation of blood samples with LPS caused an initial decrease at 3 hrs. of both DNMT1 and DNMT3a expression, followed by a progressive and significant increase in their expression **(Fig. 4).** These data suggested that the DNA methylation machinery plays an important role in controlling cytokine gene expression during infection.

**Fig 4.**
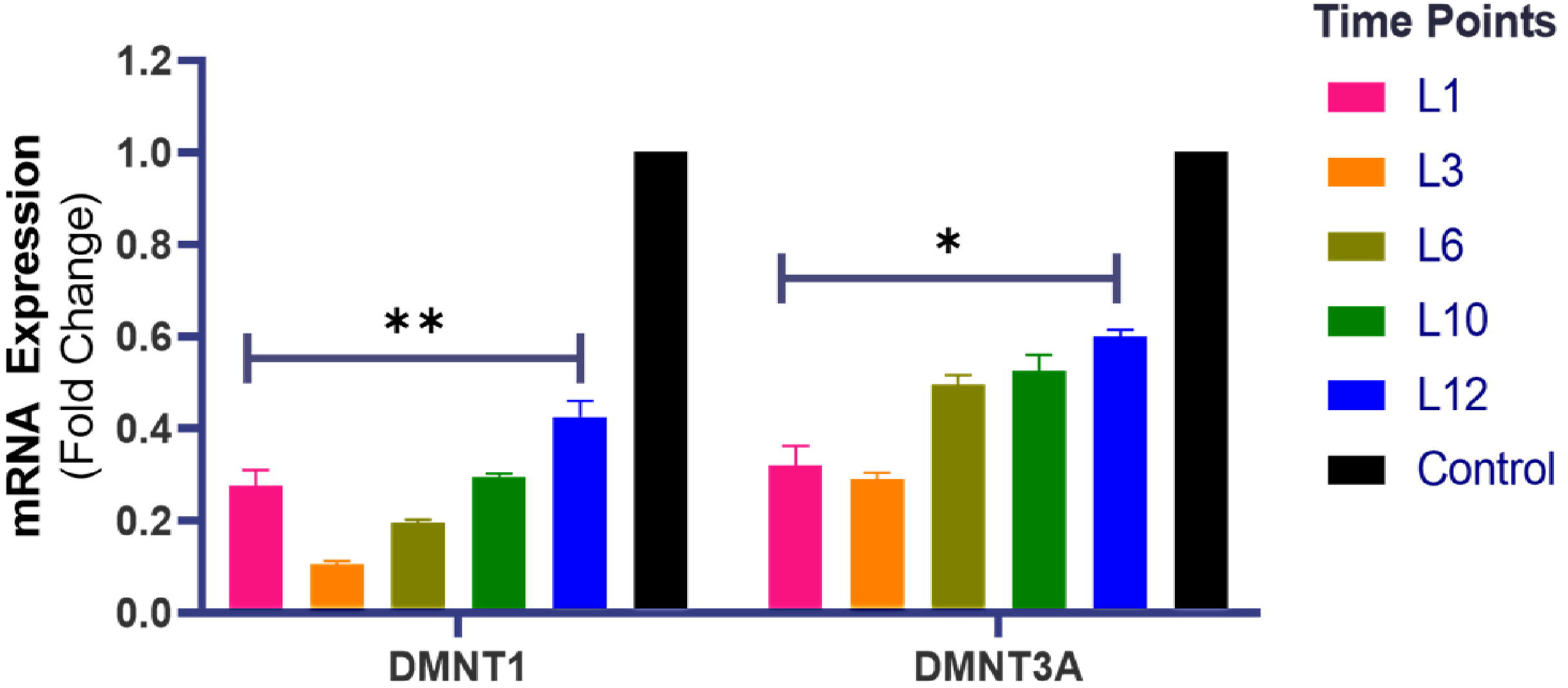
Effect of LPS treatment on DNA methyltransferases (DNMTs) expression

### DNA methylation patterns in the promoter regions of specific cytokine genes

As DNA methylation appears to play a pivotal role in regulating cytokine gene expression at the global level, changes in DNA methylation were studied for specific cytokine genes. The DNA methylation patterns in the promoter regions (+1000 bp to -100 bp) of proinflammatory (TNF-alpha, Il6, and IL1b) and anti-inflammatory (IL10) cytokine genes were investigated after whole-blood stimulation with LPS. The promoter gene sequence was retrieved from the eukaryotic promoter database (EPD), and bisulfite-specific PCR primers were designed by using software, such as Meth Primer, Zymo primer and Methyl primer. The promoter-specific PCR amplicons were purified and processed for Sanger sequencing.

The sequencing data analysis was performed using nucleotide BLAST, which is based on the principle that demethylated cytosine is converted to uracil after bisulfite conversion and is read as a thymine in subsequent PCR amplification, whereas the methylated cytosines remain unchanged. No alteration in the DNA methylation pattern in the promoter region of cytokine genes was observed following LPS treatment in the basal promoter regions of the cytokine genes in comparison to that in the untreated condition **(Fig. 5A & 5B).**

**Fig 5.**
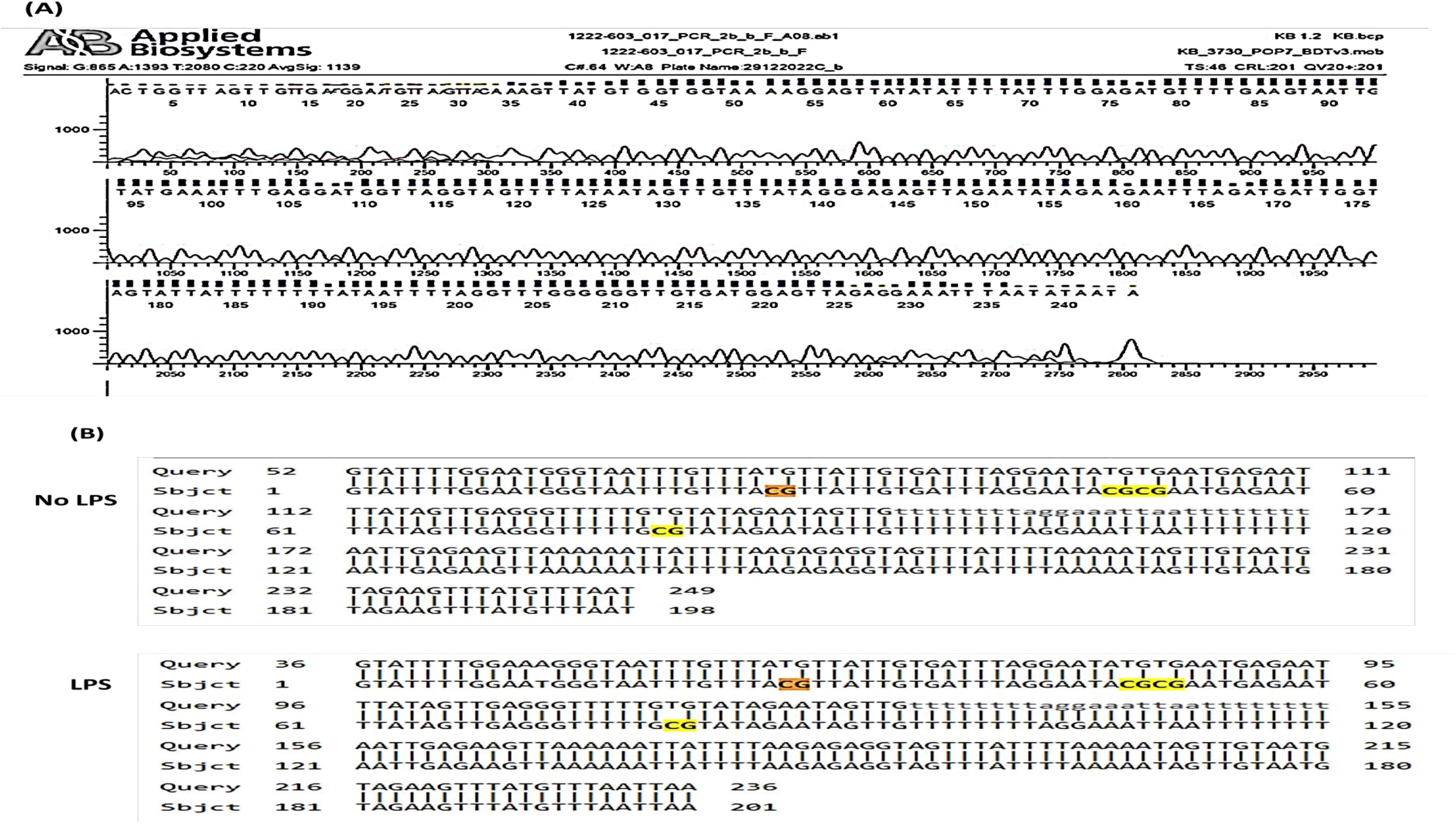
(A) Bisulfite Sanger sequencing chromatogram of the TNF-alpha promoter region. (B) Analysis of Sanger sequencing data for TNF-alpha.

### DNA methylation pattern analysis of the promoter regions of EGR1, IRF1, CREB1, c-Jun and c-Fos

As these transcription factors play pivotal roles in regulating inflammation, changes in the CpG methylation pattern in the promoter regions of these specific transcription factors were investigated via bisulfite DNA sequencing. Following LPS treatment, DNA was isolated from whole-blood samples, and the promoter regions were PCR amplified. The PCR amplicons were purified, and the bisulfited converted DNA was sequenced by Sanger sequencing **(Fig. 6A**). No change in the methylation of the EGR1 or IRF1 promoter region was observed after LPS stimulation **(Fig. 6B)**. The promoters of CREB1 and c-Fos were rapidly demethylated in response to LPS stimulation (Fig. 7A & Fig. 7B). The c-Jun promoter is methylated after LPS stimulation. These alterations in the DNA methylation patterns of the CREB1, c-Fos and c-Jun promoter regions suggested that DNA methylation and DNA demethylation play crucial roles in the rapid production of cytokine genes following endotoxin stimulation.

**Fig 6.**
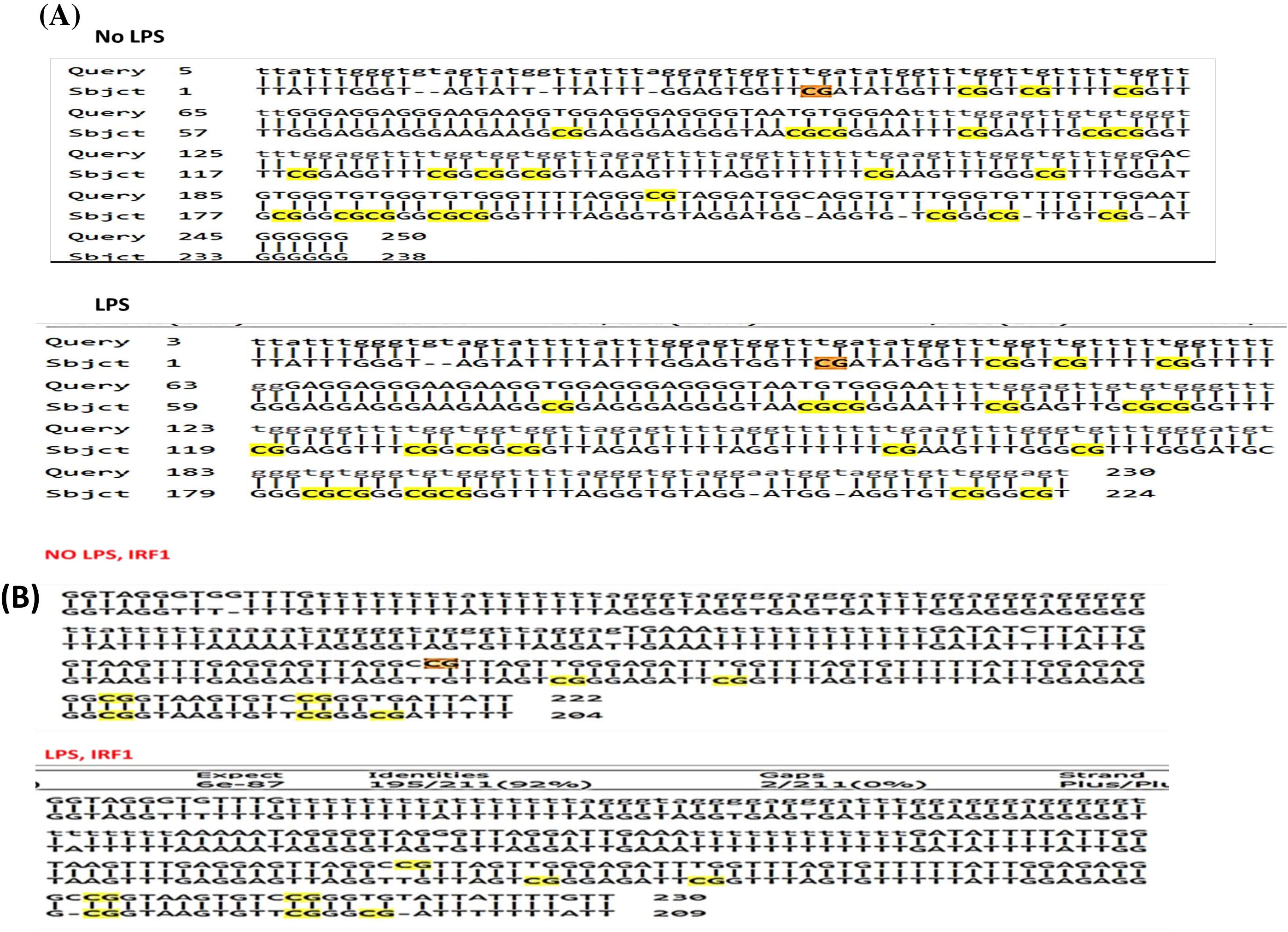
Analysis of the CpG methylation status in the promoter regions of EGR1 and IRF1 (A & B).

**Fig 7.**
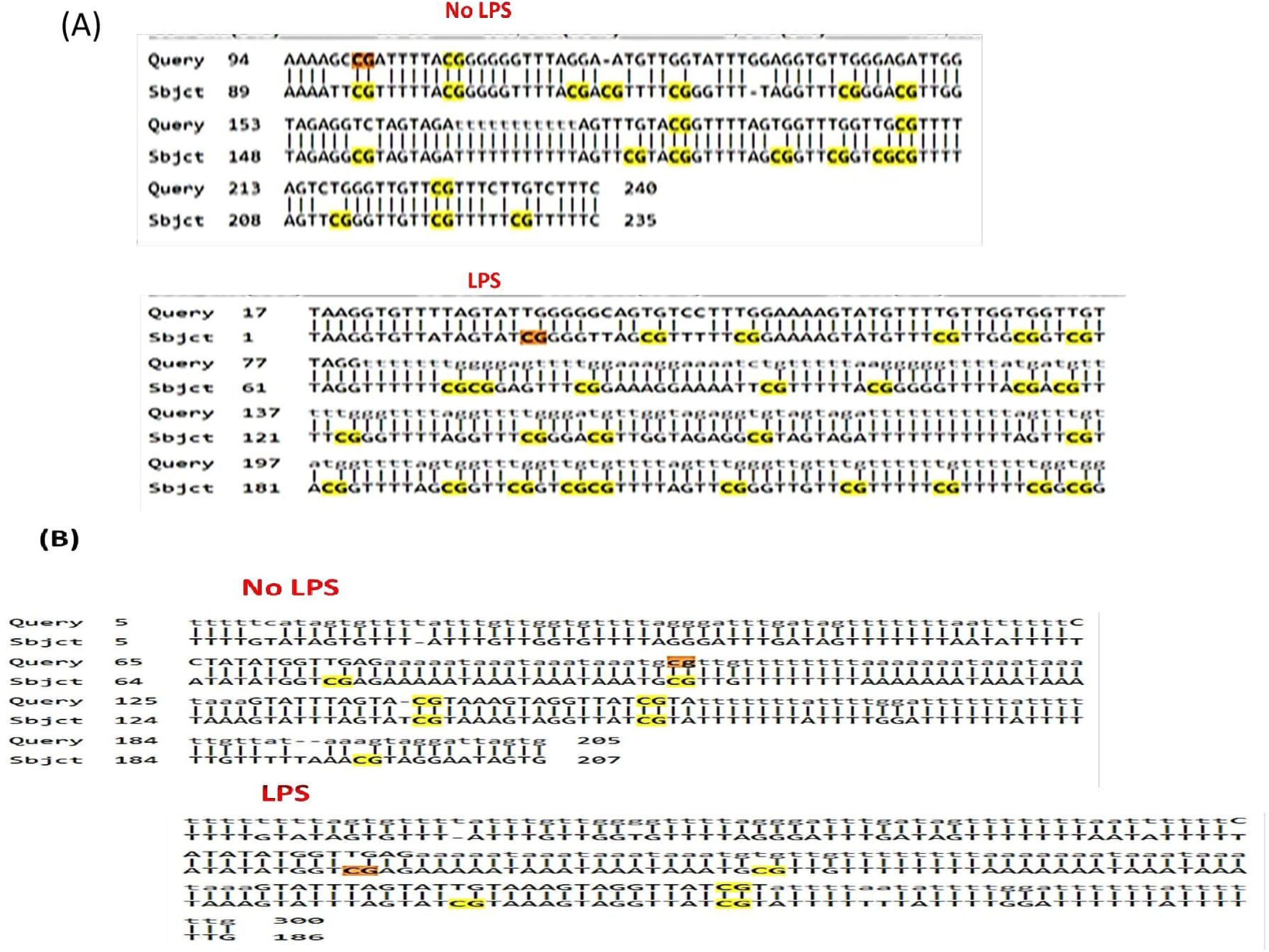
Analysis of the CpG methylation status in the promoter region of the CREB1 and c-FOS genes before and after LPS stimulation.

### Analysis of CpG methylation patterns in the promoter regions of CREB1 and c-Fos in clinical samples

Changes in the DNA methylation patterns in the promoter regions of CREB1 and c-Fos were further investigated in blood samples from pediatric cirrhotic septic and nonseptic patients. Total DNA from each blood sample was isolated, bisulfite converted and processed for Sanger sequencing **(Fig. 8A).** The CREB1 promoter region was highly demethylated in sepsis patients compared to nonsepsis patients **(Fig. 8B)**. These results suggest that CREB1 promoter DNA demethylation might play a crucial role in the regulation of cytokine gene expression in sepsis.

**Fig 8.**
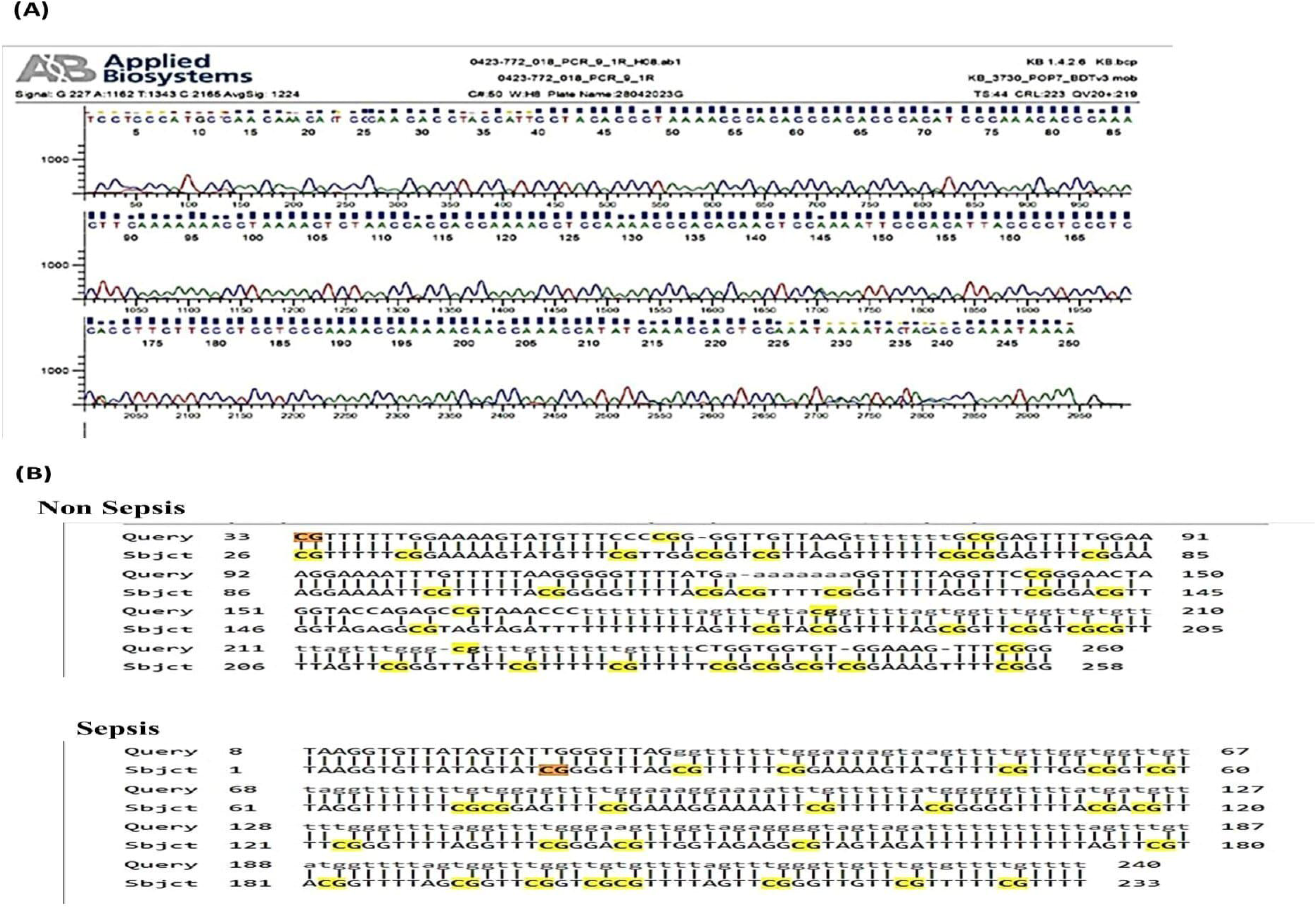
Bisulfite sequencing of pediatric patient samples. (A) Sanger sequencing chromatogram of the CREB1 promoter region. (B) Analysis of Sanger sequencing data by nucleotide BLAST. Blood samples were collected from pediatric patients (sepsis and nonsepsis). DNA was subjected to bisulfite conversion, PCR amplification, amplicon purification and Sanger sequencing. Sequencing data were analyzed by using nucleotide BLAST.

## Discussion

It is now well established that following exposure to an endotoxin at the beginning of an infection, there is a surge in the level of proinflammatory cytokines. Later, anti-inflammatory cytokines are released to neutralize their effects (Chaudhry et al., 2013). Under normal physiological conditions, cytokine release functions as a defensive mechanism. However, during uncontrolled infection, similar to sepsis, there is continuous release of proinflammatory cytokines and other inflammatory mediators, which leads to cell and tissue damage, organ failure and sometimes even death. LPS, which is a major component of gram-negative bacteria, acts as a sepsis trigger and thus plays an important role in sepsis pathogenesis. Continuous exposure to LPS activates the host immune defense mechanism to a dangerous level, as it initiates a plethora of inflammatory mediators and coagulation factors (Opal 2010, Virzi et al., 2022). The rapid increase in the expression of cytokines cannot be explained by conventional molecular changes. Hence, we propose that active DNA demethylation may play a role in the rapid overexpression of cytokine genes.

LPS stimulation of whole blood is considered to be a simple experimental model system for studying cytokines and their molecular mechanism during infection because this approach allows us to collect samples rather conveniently and requires minimal sample processing after cell culture. (Thurm et al., 2005, (Damsgaard et al., 2009, Groote et al., 1992). Accordingly, we observed that exposure to a small amount of LPS increased cytokine levels in a rather short span of time, which was sustained until 24 hours of culture. Epigenetic modifications are known to play a significant role in immune remodeling in sepsis. Histone modifications such as methylation, ubiquitination, acetylation, lactylation and DNA methylation are key signatures of gene regulation. There is evidence suggesting that histone methylation is required for the NF-κB-dependent inflammatory response. The histone demethylase JMJD3 plays a proinflammatory role in sepsis by enhancing the production of the cytokines IL6 and IL1b. During the initial stage of sepsis, the activation of histone acetyltransferases (HATs) promotes the transcription of proinflammatory genes. Additionally, histone deubiquitinating enzymes, such as USP38, play a role in innate immunity to inflammation (Holanda et al., 2021, Wu et al., 2023). Aberrant changes in global DNA methylation have also been reported in sepsis patients (Sorolla et al., 2019). However, the present study was limited to investigating the role of active DNA demethylation in cytokine gene expression. We observed that, upon LPS exposure, there was a decrease in 5mC but an increase in 5hmC within a short time span, suggesting the potential role of active DNA demethylation in sepsis. We also observed significant suppression of DNMT1 and DNMT3A expression in the presence of LPS, suggesting the potential role of DNA methylation mechanisms in sepsis. DNMT3A and TET1 act both in a competitive and collaborative manner to regulate gene expression (Gu et al., 2018). Surprisingly, we found that DNMT3A is significantly inhibited upon LPS exposure and thus contributes to the downregulation of DNA methylation during sepsis.

Surprisingly, we did not observe any alterations in the methylation patterns of the CpG sites present in the cytokine promoter regions. These results suggested that DNA methylation may not directly respond to cytokine production in response to endotoxin stimulation. Alternatively, alterations in the DNA methylation of other regions of cytokine genes could be involved in their activation. It has been reported in the literature that changes in DNA methylation at CpGs located in exons, introns and distal ends downstream of transcription start sites potentially play a role in gene expression. There are numerous transcription factors, such as CREB1, c-Fos, c-Jun, EGR1, and IRF1, that actively participate during inflammatory exposure (Smale et al., 2014). We observed that the CREB1 and c-Fos promoter regions were demethylated, whereas the c-Jun promoter region was methylated **(Fig. 5.8.2-4)**. These findings suggested that these transcription factors play pivotal roles in regulating cytokine gene expression through active DNA demethylation and methylation. c-Fos and c-Jun are components of AP-1, which is known to control inflammation and cytokine production in inflammatory skin diseases and cancers (Schonthaler et al., 2011, Qiao et al., 2016). AP-1 is known to play a role in reducing LPS-stimulated IL6 (Chen et al., 2007). The c-Fos protein interacts with the c-Jun protein to create the AP-1 complex, which controls the transcription of several proinflammatory factors. T-5224, a selective inhibitor of c-Fos/AP-1, is recognized for its substantial anti-inflammatory effects on several diseases (Zhou et al., 2022). The ETS domain of the transcription factor Elk-1 attaches to the promoter of c-FOS in response to various environmental inputs, causing an increase in its production. This results in the creation of active Fos:Jun dimers that control the activity of many Ap-1 target genes (Atsaves et al., 2019).

This study investigated the effect of DNA methylation on the regulation of cytokine genes during sepsis. Additionally, the significance of DNA methyltransferases and DNA demethylation in the context of sepsis was emphasized. The transcription factors CREB1 and c-Fos were found to be demethylated in their promoter regions during sepsis, indicating the crucial involvement of these transcription factors in sepsis. Since sepsis is a complex disease in which multiple factors contribute to its development, this work provides a more comprehensive understanding of the molecular mechanisms underlying sepsis. In addition, sepsis was shown to cause a considerable increase in the levels of numerous metabolites, which could facilitate the discovery of new biomarkers. To summarize, this study reflects a unique combination of basic and translational research. Further exploration of epigenetic mechanisms, modifiers, and omics data in sepsis could enhance the understanding of the complex pathophysiology of sepsis and facilitate the development of novel approaches for early detection and treatment.

## Supporting information

Demographic and clinical characteristics of patients enrolled in this study.PPTX

List of primers used for bisulfite sequencing.PPTX

## Ethical Approval

Ethical approval for clinical samples was obtained from the ILBS, New Delhi, India institutional human ethics committee (**IEC/2021/83/MA12)**.

## Availability of data and materials

NA

## Author contribution

AS, RK, and VK conceptualized the work. AS, PT and VG were responsible for sample processing and experimental work and were helped by JK. The data analysis was performed by AS & JK under the guidance of VK and RK. AS, RG and VK drafted the manuscript.

This manuscript has been approved by all the authors.

---

Table 1- Demographic and clinical characteristics of patients enrolled in this study

Table 2- List of primers used for bisulfite sequencing

