## Supplementary material for "DNA methylation and demethylation driven regulation of sepsis": Demographic and clinical characteristics of patients enrolled in this study.PPTX

Table 1

| Parameter | Sepsis (n=25) | Non-Sepsis (n=28) | P value |
|---|---|---|---|
| **PELD** | 25.83 (0 - 38.66) | 0 (0 - 38) | < **0.001** |
| **CTP** | 12.16 (6 - 14.66) | 7 (5 - 13) | < **0.001** |
| **AARC-ACLF** | 9.33 (5 - 13) | 5 (5 - 11) | < **0.001** |
| **Age of onset (months)** | 42.16 (2 - 162) | 114 (5 - 220) | < **0.001** |
| **Fibro scan** | 30.5 (7 - 75) | 18.65 (3.2 - 75) | < **0.001** |
| **Hb** | 9.25 (4.2 - 11.56) | 10.45 (4.3 - 15.8) | < **0.001** |
| **TLC** | 16250 (1400 - 27666.66) | 4905 (8.6 - 20200) | < **0.001** |
| **Platelets** | 159.16 (34 - 576) | 126.16 (7 - 411) | 0.02 |
| **Total Bilirubin** | 11.2 (0.8 - 29.6) | 1.5416 (0.35 - 53) | < **0.001** |
| **AST** | 186.66 (16 - 483) | 67 (21 - 1373) | < **0.001** |
| **ALT** | 103.5 (22 - 255.33) | 42.5 (11 - 533) | < **0.001** |
| **GGT** | 87.83 (24 - 800) | 39 (8 - 529) | < **0.001** |
| **Albumin** | 2.6 (1.4 - 3.6) | 3.575 (2 - 4.98) | < **0.001** |
| **Sodium** | 132 (120.66 - 140) | 138 (129 - 143) | < **0.001** |
