## Supplementary material for "DNA methylation and demethylation driven regulation of sepsis": List of primers used for bisulfite sequencing.PPTX

### Table 2

| Primer Name | Sequence | Amplicon Size(bp) |
|---|---|---|
| TNFalpha | F- GTATTTTTGATGTTTGTGTGTTTTT<br>R- TCCCTCTTAACTAATCCTCTACTAT | 287 |
| IL6 | F- GGGTAGGGTAGTAGTTAATTTTTTT<br>R- AAACTAAATTTCCTCTAACTCCATC | 265 |
| IL10 | F- GTATTTTGGAATGGGTAATTTGTTT<br>R- CTCCTTCTCTAACCTCTCTAATAAA | 298 |
| IL1b | F- GTGAGTTTATTTTTAGGGTTGTTTT R- TACACATACTTTTCTTCATTCACTT | 305 |
| IL12a | F- TAATATTTAGGTTGGGGTTTTTGTT<br>R- AAAACATTACCACACTAAATCAAAC | 275 |
| GADD45a | F- GTTTGTTTTAGTTTAAGTTGAGGTT<br>R- ACCATTAAACTATACAAATCAACCT | 266 |
| Nfkb1 | F- GTTTAAGTTTTTTTATGTGGGGAG<br>R- AACAATAATTTTAATTCACAACTCA | 279 |
| IRF1 | F- GGGTAGGTTTTTTGTTTTTTTTATTTT<br>R- CTAACTCCTAAAACCATCTTTTCAC | 291 |
| CREB1 | F- TAAGGTGTTATAGTATYGGGGTTAG<br>R-CTACRACTACCCTAACTCTACTCTTAA | 283 |
| STAT3 | F- GAGGGAGTTGTATTAGGGGTATTTA<br>R- CCACTAACCAATAAAAAAAAACAACT | 264 |
| EGF1 | F- TTATTTGGGTAGTATTTTATTTGGAGT<br>R- AAACAAAAAACCCTAATATAACAAAAC | 273 |
| KLF4 | F- TAGYGAAGGAAGTTATAAGTAAGGA<br>R- ATACCRCCAAATAAAACTAACTACC | 297 |
| FOS1 | F- TATAGTGTTTATTTGTTGGTGTTTTAG<br>R- AATTCTAACCCTAATTCAATACAAAAC | 259 |
| JUN1 | F- GTAGTYGTGAATTTGAGTTTTTTTT<br>R- CATAAAAACTCCACCCTAAAAAATT | 285 |
